# Transposon insertion causes *ctnnb2* transcript instability that results in the maternal effect zebrafish *ichabod* (*ich*) mutation

**DOI:** 10.1101/2025.02.28.640854

**Authors:** Zsombor Varga, Ferenc Kagan, Shingo Maegawa, Ágnes Nagy, Javan Okendo, Shawn M. Burgess, Eric S. Weinberg, Máté Varga

**Affiliations:** Department of Genetics, ELTE Eötvös Loránd University, Budapest, Hungary; Max Planck Institute for Multidisciplinary Sciences, Göttingen, Germany; Department of Intelligence Science and Technology, Graduate School of Informatics, Kyoto University, Japan; Hungarian Defence Forces Medical Centre, Budapest, Hungary; Translational and Functional Genomics Branch, National Human Genome Research Institute, Bethesda, MD, USA; Department of Biology, University of Pennsylvania, Philadelphia, PA, USA

## Abstract

The maternal-effect mutation *ichabod* (*ich*) results in ventralized zebrafish embryos due to impaired induction of the dorsal canonical Wnt-signaling pathway. While previous studies linked the phenotype to reduced *ctnnb2* transcript levels, the causative mutation remained unidentified. Using long-read sequencing, we discovered that the *ich* phenotype stems from the insertion of a non-autonomous CMC-Enhancer/Suppressor-mutator (CMC-EnSpm) transposon in the 3’UTR of the gene. Through reporter assays, we demonstrate that while wild type *ctnnb2* mRNAs exhibit remarkably high stability throughout the early stages of development, the insertion of the transposon dramatically reduces transcript stability. Genome-wide mapping of the CMC-EnSpm transposons across multiple zebrafish strains also indicated ongoing transposition activity in the zebrafish genome. Our findings not only resolve the molecular basis of the *ich* mutation but also highlight the continuing mutagenic potential of endogenous transposons and reveal unexpected aspects of maternal transcript regulation during early zebrafish development.

## Introduction

In sexually reproducing species, new life is initiated by the fusion of parental gametes, albeit the zygote is anything but a blank slate after fertilization. Oocytes are loaded with maternal factors, mRNAs and proteins, which drive the developmental processes during the earliest stages of embryogenesis, until transcription from the embryo’s own genome is turned on. The length of this stage and the timing of the so called maternal-zygotic-transition (MZT), which effectively terminates this early stage of development, is highly species-dependent (Vastenhouw *et al*, 2019; Brantley & Talia, 2024). In those species where several cell cycles elapse before MZT, usually the oocytes are already highly patterned themselves. Consequently, the maternal factors will not only coordinate early cell divisions but will also have a significant role in the initial patterning of the embryonic axes (Fuentes *et al*, 2020; Abrams & Mullins, 2009).

The importance of maternal genes was established during the initial dissection of the early metazoan development using forward genetics in fruit flies (*Drosophila melanogaster*) (reviewed in (Johnston & Nüsslein-Volhard, 1992; Wieschaus & Nüsslein-Volhard, 2015). Subsequent genetic screens in zebrafish (*Danio rerio*), recently complemented by “crispant” screens, have also identified numerous important maternal effect genes (Pelegri & Schulte-Merker, 1999; Dosch *et al*, 2004; Wagner *et al*, 2004; Pelegri *et al*, 2004; Moravec *et al*, 2021). Some maternal mutants, such as *ich* (Kelly *et al*, 2000) or *tokkaebi* (*tkk*) (Nojima *et al*, 2004, Nojima *et al*, 2010) were also uncovered independently of forward mutagenesis.

The mapping and characterization of these maternal mutants (in parallel with other experiments) revealed the intricate relationship between the maternal factors asymmetrically segregated along the animal-vegetal axis and the dorsal-specific activation of the canonical Wnt/β-catenin signaling pathway, which occurs in early blastula stages (Langdon & Mullins, 2011; Fuentes *et al*, 2020). Fertilization of the zebrafish egg is followed by a cortical rotation, which will move the vegetally located dorsal determinant (likely *wnt8a* mRNA) to the future dorsal side of the developing embryo (Hibi *et al*, 2018; Tran *et al*, 2012; Ge *et al*, 2014; Lu *et al*, 2011). This relocation is essential for the timely activation of the canonical Wnt pathway and the nuclear localization of β-catenin (Ge *et al*, 2014; Tran *et al*, 2012).

The maternal-effect *ich* mutation results in ventralized embryos, due to impaired induction of dorsal canonical Wnt-signaling, resulting from reduced *ctnnb2* transcript levels (Bellipanni *et al*, 2006; Kelly *et al*, 2000). This property makes it particularly well suited to study early dorsal patterning events in development (Tsang *et al*, 2004; Maegawa *et al*, 2006; Varga *et al*, 2007; Cruz *et al*, 2010; Varga *et al*, 2011; Tanaka *et al*, 2017; Wu *et al*, 2012). Of note, the mutation can be completely rescued by the injection of *ctnnb2* mRNA, *ctnnb1* mRNA or *Xenopus* β-catenin mRNA as well (Kelly *et al*, 2000; Bellipanni *et al*, 2006).

While genetic mapping of the *ich* allele suggested that the mutation resides in the proximity to *ctnnb2*, no mutation was found in the coding sequence (CDS) of the gene in *ich* homozygotes. Furthermore, the telomeric location of the *ctnnb2* locus frustrated previous attempts to positionally clone the mutation (Bellipanni *et al*, 2006).

Ever since their discovery in the middle of the 20^th^ century (McClintock, 1950), transposable elements (TEs) have been recognized as potent biological mutagens capable of disrupting the normal function of genes in a variety of organisms (Hayward & Gilbert, 2022). Indeed, some historically relevant phenotypes, such as the wrinkled peas of Mendel (Bhattacharyya *et al*, 1990) or the melanized forms of the peppered moths which appeared during the Industrial Revolution in 19^th^ century Britain (Hof *et al*, 2016) were the result of transposon insertions. The mobility of these “jumping genes” have also made them a prime tool for transgenic technologies in multiple model species, including zebrafish (Kawakami *et al*, 2017; Ivics *et al*, 2009; Kwan *et al*, 2007; Korzh, 2007; Kawakami, 2007; Varshney & Burgess, 2014).

Vertebrate genomes carry varying levels of TEs, and in some species such as zebrafish and humans over half of the genome can consist of such sequences (Howe *et al*, 2013; Chalopin *et al*, 2015). Type II DNA transposons are in general enriched within the zebrafish genome and CMC EnSpm transposons constitute one of the largest family amongst them (Howe *et al*, 2013; Gao *et al*, 2016; Chalopin *et al*, 2015; Shao *et al*, 2019).

Using a long-read genomic sequencing approach, here we show that the *ich* phenotype results from the insertion of an EnSpm-type transposon into the 3’UTR of the *ctnnb2* gene, which causes the degradation of an otherwise very stable maternal transcript.

## Results

### A transposon insertion can be observed in the 3’UTR of *ctnnb2* in *ich* mutants

Earlier positional cloning experiments placed the causative mutation of *ich* in close proximity to the *ctnnb2* locus, and while no mutations could be uncovered in the CDS of the gene, maternal levels of *ctnnb2* mRNA and Ctnnb2 protein were severely reduced in the embryos of homozygous females (Bellipanni *et al*, 2006).

To determine the exact physical nature of the causative mutation, we conducted Oxford Nanopore Technologies (ONT) long-read sequencing on one homozygous *ich* female. We have assembled the ONT reads into contigs and aligned the resulting contigs to a newly assembled, “telomere-to-telomere” sequence of chromosome 19 (GCA_033170195.2) and selected the contig that spanned the *ctnnb2* locus (Figure 1A,B). The alignment of this 2.27 Mbp long contig to the reference genome sequence showed no sign of structural variation, thus proceeded to a manual annotation of the *ctnnb2* locus. This annotation revealed the presence of a 2469 bp long insertion in the last exon of the gene, within the 3’UTR region (Figure 1C).

**Figure 1:**
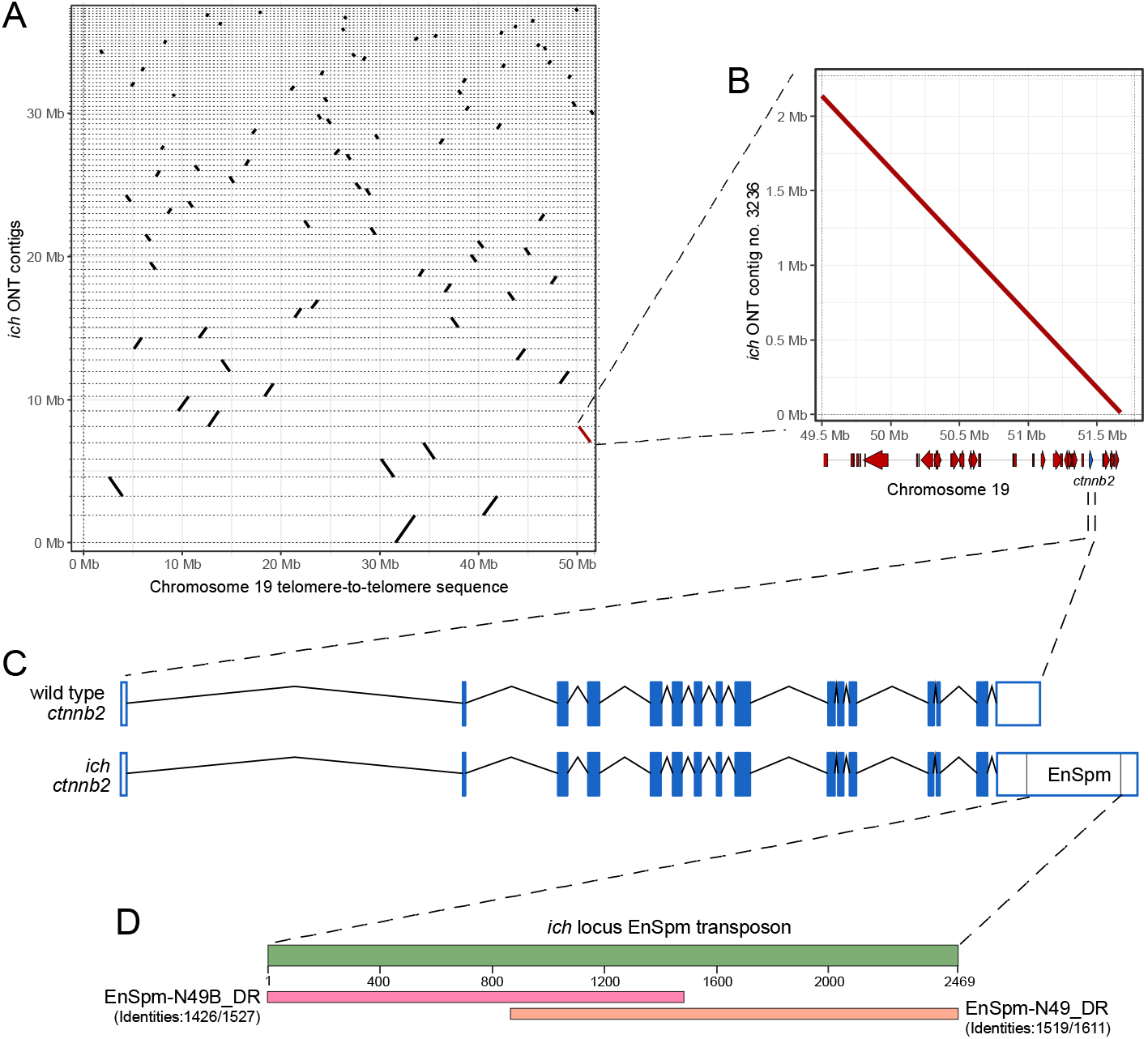
Long-read sequencing identifies an EnSpm transposon in the 3’UTR of *ctnnb2* in *ich* animals. (A) Dotplot representation of different *ich* contigs that could be aligned to the “telomere-to-telomere” sequence of the zebrafish chromosome 19. The highlighted (red) contig contains the genomic region of the *ctnnb2* gene. (B) Dotplot comparison of the relevant *ich* contig and the telomeric portion of chromosome 19 reveals no genomic rearrangement in this region in *ich* animals. Gene position schematics are visible along the horizontal axis. (C) Comparison of wild-type and *ich*-specific *ctnnb2* sequences showing the position of the EnSpm-type transposon insertion. (D) BLAST results of the *ich*-specific transposon in the FishTEDB (Shao *et al*, 2024).

A closer examination of this insertion revealed that it is a hybrid sequence between two previously recognized, non-autonomous CMC-EnSpm type transposons: EnSpm-N49_DR and EnSpm-N49B_DR (Figure 1D) (Shao *et al*, 2024).

With allele-specific primers, we were also able to confirm that in all *ich* homozygous females both *ctnnb2* alleles carried this insertion (Supplementary Figure 1).

### *ctnnb2* transcripts show high stability during early development

Previous work has already highlighted that while the *ich* mutation causes a reduction in the level of *ctnnb2* transcripts, the phenotype itself can be rescued not only by the injection of *ctnnb2* mRNA, but also *ctnnb1* mRNA can rescue the phenotype (Bellipanni *et al*, 2006). Zebrafish Ctnnb1 and Ctnnb2 are highly similar proteins, with slight differences detected only in the C-terminal region (Supplementary Figure 2A,B). The transcripts of the two orthologs also show high similarity within their coding regions and 5’UTRs (83.11% identity). Sequence comparisons, however, also show that the 3’UTR of the two genes is highly divergent, with no similarities to be found (Supplementary Figure 2C). As both paralogs are also transcribed during oogenesis (Supplementary Figure 3D), the difference in the 3’UTRs suggested that one plausible explanation for the inability of endogenous *ctnnb1* to rescue *ich* would be a difference in the stability of *ctnnb1* and *ctnnb2* maternal transcripts.

The lack of adequate labeling methods and paralog-specific antibodies have also limited the ability to discern the dynamics of maternal and zygotic transcripts and protein products in earlier experiments. Newly available datasets that rely on metabolic labeling to differentiate between maternal and zygotic transcripts and assess maternal protein stability (Bhat *et al*, 2023; Pescador *et al*, 2024; Fishman *et al*, 2024), however, gave us new opportunities to compare the expression of *ctnnb1* and *ctnnb2* at high resolution during the early development of wild-type embryos.

Interestingly, we observed that *ctnnb2* transcripts are about three times as abundant as *ctnnb1* ones both pre- and post-MZT (Figure 2A), even though no zygotic transcription can be detected from the former gene (Figure 2B). Similarly, Ctnnb2 levels also seem constant during these early stages of development (Figure 2C). Of note, while protein profiling datasets did not report Ctnnb1 stability (Pescador *et al*, 2024), earlier observations using a pan-β-catenin antibody showed that maternal Ctnnb1 is present in both wild-type and *ich* embryos (Bellipanni *et al*, 2006; Valenti *et al*, 2015).

**Figure 2:**
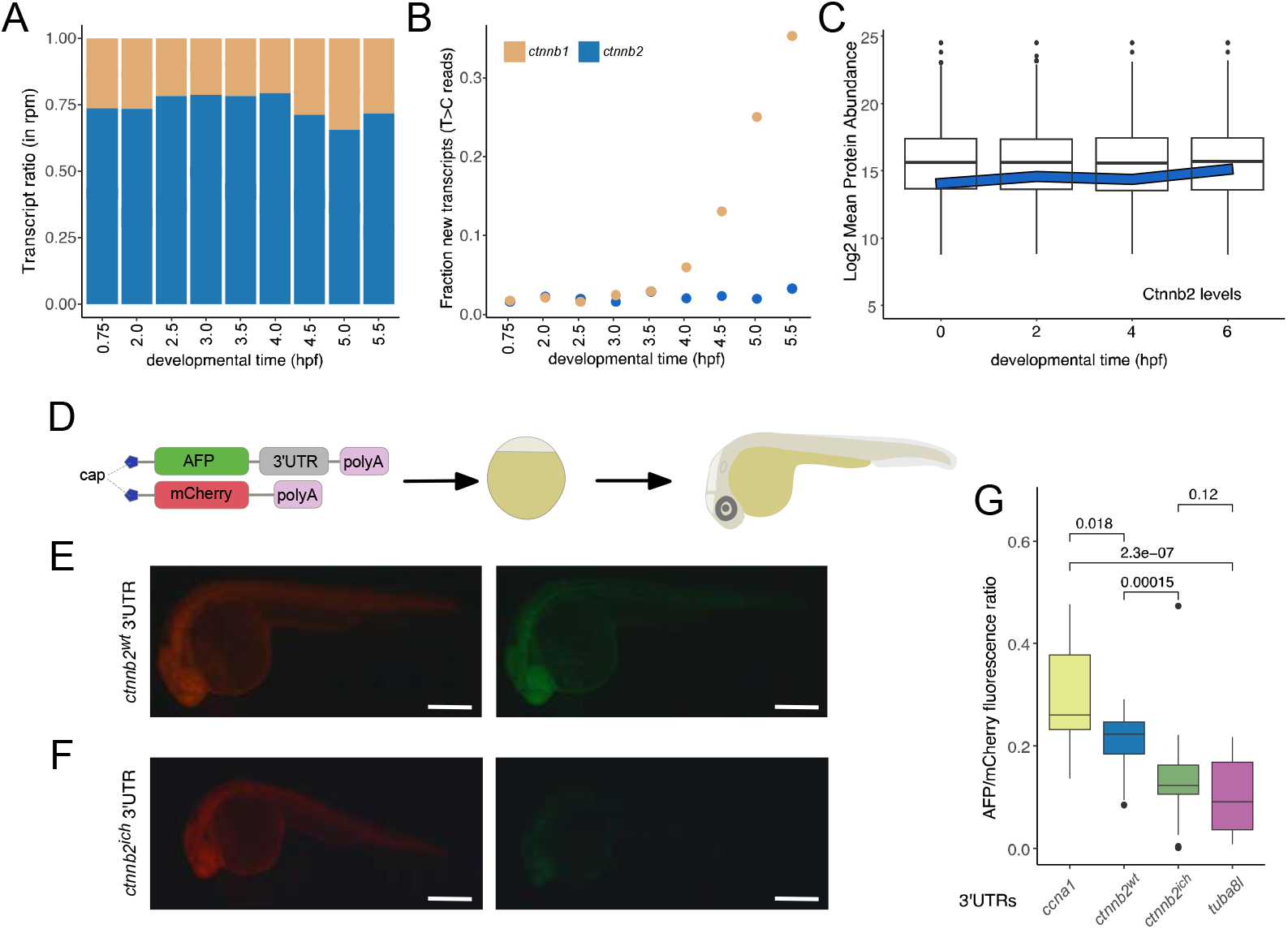
High stability of maternal *ctnnb2* transcripts is disrupted by 3’UTR transposon insertion. Transcriptome analysis of maternal *ctnnb1* and *ctnnb2* mRNAs shows a heightened abundance of maternal *ctnnb2* transcripts throughout early development (rpm – reads per million bp). (B) Metabolic labelling suggests active transcription of *ctnnb1* after *ZGA*, but no expression of *ctnnb2*. (Data for panels A and B are derived from (Bhat *et al*, 2023). (C) Maternal Ctnnb2 protein levels are also very stable during early stages of development as compared to mean protein abundances at these timpoints. (Data from (Pescador *et al*, 2024). (D) Experimental scheme used to test the effect of 3’UTRs on the stability of mRNAs. *In vitro* transcribed *AFP* mRNAs with different 3’UTRs are combined with *mCherry* transcripts with invariant 3’UTRs and injected into 1-cell stage zebrafish embryos. Fluorescence intensity for both AFP and mCherry is measured at 24 hpf. (E, F) Typical fluorescence observed in embryos injected with *AFP* carrying *ctnnb2*^*wt*^ or *ctnnb2*^*ich*^ 3’UTRs, respectively. (Scale bars: 0.5 mm) (G) Relative fluorescence intensities observed in embryos injected with different 3’UTRs compared to stable and unstable controls (*ccna1* and *tuba8l* 3’UTRs, respectively).

### Transposon insertion in the 3’UTR disrupts the stability of *ctnnb2* transcripts

To test if the insertion of the hybrid EnSpm transposon in the 3’UTR destabilizes the *ctnnb2* we utilized a previously described reporter assay (Giraldez *et al*, 2006). We co-injected *in vitro* synthesized *AFP* mRNAs with varying 3’UTRs and *mCherry* with non-variant 3’UTRs in wild type embryos and estimated the ratio of AFP and mCherry intensity in these embryos at 24 hpf (Figure 2D).

We measured intensities for constructs carrying either the wild-type or the *ich*-specific *ctnnb2* 3’UTRs and compared them to the well-established stable and unstable 3’UTRs of *ccna1* and *tuba8l*, respectively (Giraldez *et al*, 2006). These results show that while the stability of wild type *ctnnb2* 3’UTR is comparable to that of the extremely stable *ccna1* 3’UTR, the insertion of the EnSpm transposon resulted in a dramatic reduction in the stability of the transcript, making it similar to that observed for *tuba8l*.

The 3’UTR of the wild-type *ctnnb2* gene is relatively long and carries a number of secondary structural motifs (Supplementary Figure 3A), that could either stabilize the transcript by themselves, or could serve as platforms for interaction with RNA-binding proteins (RBPs) that regulate the stability of the mRNA (Ho *et al*, 2021). To reveal if the insertion of the transposon could introduce any destabilizing sequence motifs in the 3’UTR, we took advantage of a random forest model to compare the predicted stabilities of the different *ctnnb2* 3’UTR sequences with the *ctnnb1* 3’UTR (Vejnar *et al*, 2019). Interestingly, this analysis did not reveal any destabilizing motifs within the transposon itself, therefore the observed decrease in transcript stability is most likely not due to the potential destabilizing effects of particular sequences within the transposon itself (Supplementary Figure 3B).

Active transposons are often the target of piRNA-driven destruction during zebrafish germ-cell development and embryogenesis (Houwing *et al*, 2007a, 2008; Wei *et al*, 2012; Kaaij *et al*, 2013), which offers another possibility for the premature degradation of maternal *ctnnb2* mRNAs. To test if this could be a feasible explanation for *ich* embryos, first we looked at the temporal dynamics of EnSpm-N49/N49B transcription. The reanalysis of previously published transposon expression datasets (Chang *et al*, 2022) shows that these particular transposons become active around the time of the zygotic genome activation (ZGA) but become downregulated shortly after. The decrease in the levels of EnSpm-N49/N49B transcripts around shield stage is concomitant with an increase in the amount of piRNAs (Wei *et al*, 2012), suggesting a Piwi-dependent silencing mechanism, which could also account for the degradation of the *ich-*specific *ctnnb2* transcript as well.

### Active EnSpm-type elements in the zebrafish genome

Our mapping results revealed the recent disruption of the *ctnnb2* 3’UTR in *ich* animals due to the transposition of a hybrid EnSpm-N49/N49B-type transposon. As this suggested that the transposon is still active, we decided to map the location of EnSpm-N49, EnSpm-N49B and EnSpm-N49/49B transposons in the zebrafish reference genome and compare it to other, recently sequenced wild-type strains.

Our results suggest that of the three transposons the hybrid one appears to be the most active, as we were able to detect 72 copies in the GRCz11 reference genome, whereas only 7 copies of EnSpm-N49 and 16 copies of EnSpm-N49B were detected using an 80% sequence similarity threshold (Figure 3A, Supplementary Figure 4A). Our analysis also shows higher sequence similarity for EnSpm-N49/N49B copies than for the other transposons, suggesting that this particular transposon is evolutionarily novel (Supplementary Figure 4A). The genomic distribution of all three transposons is relatively similar, with most copies found in distal intergenic or intronic regions (Supplementary Figure 4B-D).

**Figure 3:**
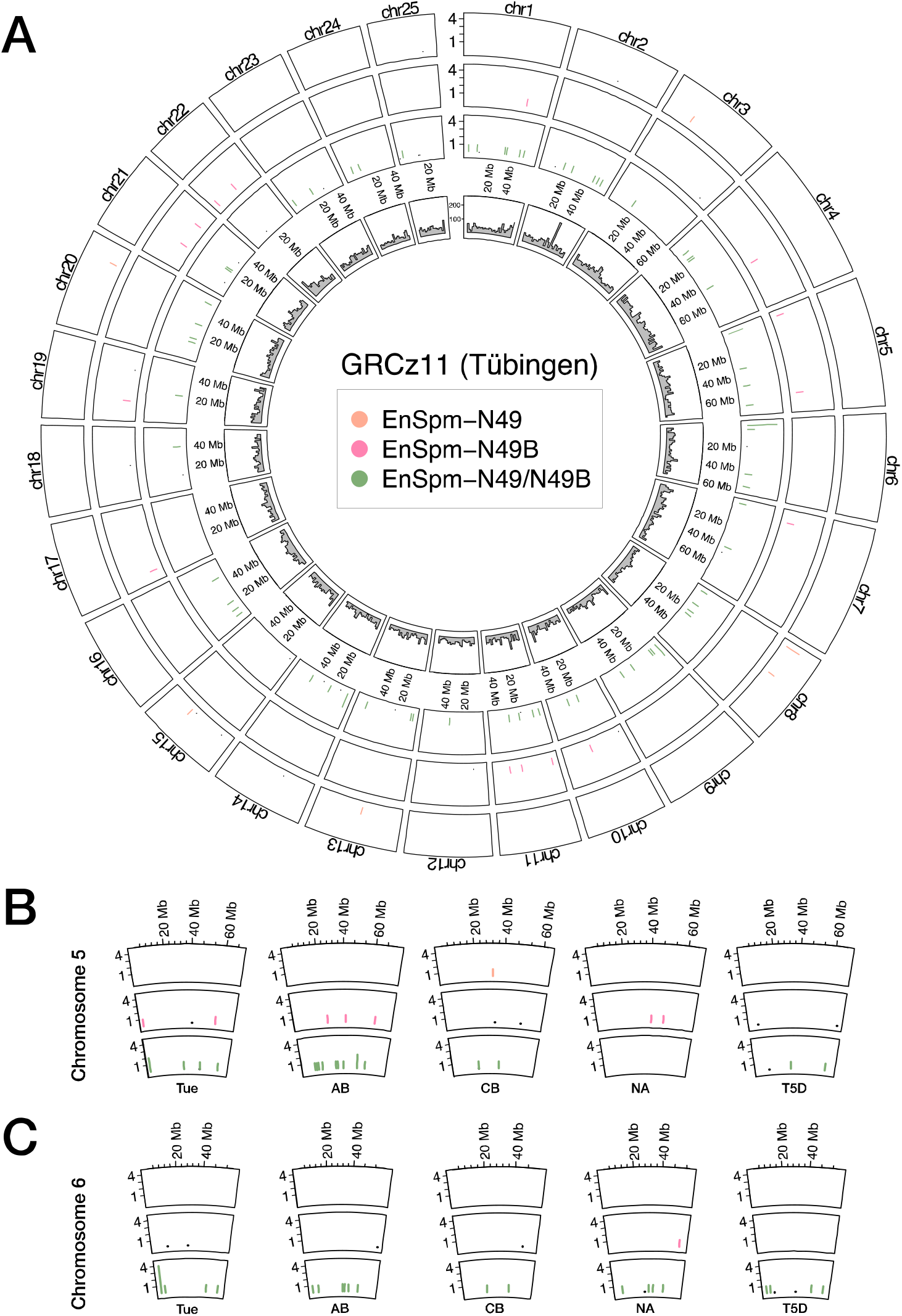
The position of EnSpm transposons observed in this study vary across zebrafish genomes. Genomic positions of EnSpm-N49 (orange), EnSpm-N49B (pink) and EnSpm-N49/N49B (green) transposons in nonoverlapping 2-Mbp windows across the nuclear chromosomes of the Tübingen (Tue) GRCz11 reference genome. Inner circle shows the coverage of protein-coding genes on the same chromosomes. (B, C) Genomic positions of the observed transposons show variability across the genomes of multiple wild-type isolates for chromosomes 5 (B) and 6 (C). (Y axis denotes copy numbers in the 2-Mbp windows. For other chromosomes see Supplementary Figure 4.)

To compare the positions of the investigated transposons between different wild-type strains, we also determined the location of the transposons in AB, Cooch Behar (CB), Nadia (NA) and T5D strains (Balik-Meisner *et al*, 2018; Deng *et al*, 2022). Our results confirm that the hybrid EnSpm-N49/N49B appears to be the most active of the three transposons (Supplementary Figure 4E-H), and also reveal that all three transposons are actively transposing in the zebrafish genome as they show divergent distributions in the different isolates (Figure 3B,C).

## Discussion

The mobility and hence the mutagenic potential of mobile genetic elements has been obvious since their discovery in maize (McClintock, 1950) and it is now recognized as a general feature all living things (Hayward & Gilbert, 2022). The molecular characterization and in-depth understanding of the transposition process has also made it possible to use such “selfish genetic elements” in biotechnology and numerous transgenesis methodologies have been developed that exploit the mobility of these sequences essentially in all model organisms (Amsterdam *et al*, 2004; Kwan *et al*, 2007; Korzh, 2007; Kawakami, 2007; Ivics *et al*, 2009; Varshney & Burgess, 2014; Kawakami *et al*, 2017). Thanks to the related expansion of the genetic toolbox over the years numerous zebrafish mutant lines have been identified resulting from retroviral- or transposon-based mutagenesis screens (Amsterdam *et al*, 2004; Nagayoshi *et al*, 2007; Kotani & Kawakami, 2008).

Besides exogenous mobile elements, however, endogenous ones can also have mutagenic effects. Indeed, over the past decades several zebrafish mutant alleles known to be caused by spontaneous insertion of endogenous transposons have been also identified (Streisinger *et al*, 1986; Nojima *et al*, 2010). The sudden expansion of type II DNA transposons, in general, and CMC EnSpm transposons, in particular, in the zebrafish genome has expanded the repertoire of putative biological mutagens in this species (Howe *et al*, 2013; Gao *et al*, 2016; Chalopin *et al*, 2015; Shao *et al*, 2019). As shown by our results characterizing the *ich* allele, at least some of these transposons are still active (Figure 3, Supplementary Figure 4) and exert a mutagenic influence. While TEs are often enriched on chromosome 4, the ancestral sex-chromosome of zebrafish with an abundance of transposable elements (Wilson *et al*, 2014; Chang *et al*, 2022; Wilson & Postlethwait, 2024), and the accumulation of CMC EnSpm transposons has been also associated with the evolution of sex chromosomes in some other species (Schemberger *et al*, 2019), we do not see an enrichment of the three observed transposons on chromosome 4.

The destabilization of *ctnnb2* transcripts due to the insertion of a EnSpm-N49/N49B transposon sequence in exon 16 of the gene, encoding the 3’UTR (Figure 1 and 2) also highlights that despite years of rapid advances, our understanding of the processes that regulate the stability and turnover of maternal mRNAs is still incomplete. Previous work has already highlighted that the presence of miR-430 binding sites (Giraldez *et al*, 2006; Liu *et al*, 2020), the codon composition of the CDS (Bazzini *et al*, 2016; Medina-Muñoz *et al*, 2021), the presence of specific structural elements in the transcript 3’UTRs (Mishima & Tomari, 2016; Vejnar *et al*, 2019) as well as the length of the polyA tail (Subtelny *et al*, 2014) can all have essential roles in regulating mRNA stability during MZT. As the transposon insertion does not appear to introduce destabilizing structures into the 3’UTR (Supplementary Figure 3), in the case of the *ctnnb2*^*ich*^ allele the likeliest explanation involves the piRNA-dependent degradation of the mRNA.

The Piwi-piRNA pathway provides a small RNA-based adaptive defence against TEs in most eukaryotic genomes (Aravin *et al*, 2007; Czech *et al*, 2018; Ozata *et al*, 2019; Iwasaki *et al*, 2025). It has especially important roles in containing these biological mutagens during the development of the gametes in most animals and zebrafish is no exception (Houwing *et al*, 2007b, 2008; Zhou *et al*, 2010; Kaaij *et al*, 2013). Elevated activity of the Piwi-piRNA pathway during oogenesis is also the most likely cause for the degradation of the *ctnnb2*^*ich*^ transcripts, which contain a perfect transposon target.

It is also worth highlighting the high stability of the wild-type maternal *ctnnb2* transcript. Even in the absence of a zygotic component, there seem to be approximately three times as many *ctnnb2* transcripts as those of *ctnnb1* during early development (Figure 2). This does not appear to be caused by differential stability of 3’UTRs as based on this sequence alone, *ctnnb1* transcripts should be at least as stable as their *ctnnb2* counterparts (Supplementary Figure 3).

This makes it likely that an interaction with yet unknown RNA-binding proteins (RBPs) is behind the observed stability levels. It is noteworthy, however, that a screen of putative RNA binding sites (Benoit Bouvrette *et al*, 2019) did not highlight any obvious candidates that would account for the differential stability of *ctnnb1* and *ctnnb2* (not shown). Understanding the causes for the high stability of *ctnnb2* transcripts, therefore, could provide important additional information about the mechanisms that preserve transcripts during oogenesis and protect them from degradation during ZGA, when typically, we see the turnover of maternal transcripts to zygotic ones.

In summary we have identified the insertion of an EnSpm transposon in the 3’UTR of the *ctnnb2* gene as the most likely proximate cause of *ctnnb2* transcript instability observed in the embryos of homozygous *ich* mutant fish. We showed that the transposon-containing *ctnnb2* 3’UTR is considerably less stable than the wild-type counterpart. We also describe the genomic distribution of the transposon in multiple zebrafish genomes, providing evidence about the recent activity of this TE.

## Materials and Methods

### Fish husbandry and maintenance

Adult wild-type (*ekwill*) and *ich* (*ctnnb2*^*p1*^) fish used for these experiments were maintained and bred in the animal facility of the Biology Institute of ELTE Eötvös Loránd University according to existing protocols (Aleström *et al*, 2019; Westerfield, 2000). All animal husbandry protocols used for this study were approved by the ELTE Animal Welfare Animal Committee and the Hungarian National Food Chain Safety Office and by the Animal Experiment Committee of Kyoto University.

### Genomic DNA isolation and long-read sequencing

Total genomic DNA was isolated from of a homozygous *ich* female specimen (weighing approximately 50 mg) stored in DNA/RNA Shield Reagent (Zymo Research, cat # R1100) using a DNeasy Blood and Tissue Kit (Qiagen, cat # 69504) in accordance with the manufacturer’s protocol for Purification of Total DNA from Animal Tissues, with some modifications. In summary, 360 µl of ATL buffer and 40 µl of proteinase K were added to the fish sample, which was then vortexed rigorously. The mixture was then incubated at 56°C for 3.5 hours with 400 rpm mixing and vortexing every 30 minutes. Thereafter, 400 µl of AL buffer and 400 µl of ethanol were added to the lysate, which was then vortexed vigorously. The subsequent steps were then continued according to the manufacturer’s protocol. Two sequencing libraries were prepared with the Ligation Sequencing Kit SQK-LSK110 (Oxford Nanopore Technologies, Oxford, cat # SQK-LSK110) according to the manufacturer’s protocol for genomic DNA by ligation, with 1.5 μg of input from both genomic DNAs being added to each library preparation. The entire quantity of each prepared library (1^st^ library: 680 ng, 2^nd^ library: 39.6 ng) was loaded in 75 μl volume onto two R9.4.1 flow cells (Oxford Nanopore Technologies, cat # FLO-MIN106) and run for 48 hours until flow cell extinction. Sequencing data were generated using the Oxford Nanopore GridION platform and performing real-time super-accurate basecalling with MinKNOW 23.04.5 and Guppy 6.5.7 (Oxford Nanopore Technologies, Oxford, UK) to achieve the highest possible accuracy rate. The sequencing runs generated a total of 28.19 Gb of raw genomic data in two batches.

### Genome assembly and analysis

Raw ONT data was uploaded into the https://usegalaxy.euplatform (Community *et al*, 2024). For assembly of contigs we used the *Fly* assembler (Kolmogorov *et al*, 2019), and later performed scaffolding based on the GRCz11 reference genome sequence using *RagTag* (Alonge *et al*, 2022). This scaffolding step allowed us to select for those *ich* contigs that contained genomic sequences specific for chromosome 19. These sequences were aligned along the chromosome-to-chromosome sequence of chromosome 19 using *minimap2* (Li, 2018, 2021). The settings used to run these programs are presented in Table 1. The contig containing the genomic neighborhood (contig no. 3236) was annotated manually in the Benchling online software suite (Benchling [Biology Software], 2025).

**Table 1:** Standard parameters used in the usegalaxy.eu pipeline.

| Software package | Parameters used |
| --- | --- |
| Fly | --nano-raw ./input_0.fastq.gz -o out_dir -t -i 1 |
| RagTag | scaffold -u --aligner minimap2 --mm2-params '-k19 -w19 -A1 -B19 -O39,81 -E3,1 -s200 -z200 --min-occ-floor=100' -f 1000 --remove-small -q 10 -d 100000 -i 0.2 -a 0.0 -s 0.0 -r -g '100' -m '100000' -o ./ -t |
| minimap2 | -x asm5 --q-occ-frac 0.01 -t |

### Genomic DNA isolation for genotyping PCRs

PCR-ready genomic DNA isolation was performed as described before (Meeker *et al*, 2007). Briefly, after anesthesia, each fish was fin-clipped and tissue samples were transferred to a 200 μL PCR tube. After the addition of 100 μL of 50 mM NaOH solution the samples were heated at 95°C for 15 minutes with occasional flicking. Subsequently, the tubes were cooled down to 4°C and lastly 10 μL of 1 M Tris-HCl (pH 8.0) was added for balancing the pH to near-neutral.

### Genotyping PCR of *ich* fish

Genotyping PCRs were performed using with the following three primer pairs: (1) the ctnnb2_ex16 primer pair, ctnnb2_ex16 L (5’ – TCCGTGTTCCCAGAAGAAGC – 3’) and ctnnb2_ex16 R (5’ – GAAAGTGCCTGATGAGTGCG – 3’), (2) the ctnnb2_ex16_tp primer pair, ctnnb2 ex16 tp L (5’ – CTAGGATCCGTTTGTCAAGCCCACACACC – 3’) and ctnnb2_ex16_tp R (5’ – GATGAATTCCTCATGGTACTGACCCGAGC – 3’), and (3) ctnnb2_ex16_whole primer pair, ctnnb2_ex16_whole L (5’ – GCGTGTGGTTAATCGTCTGC – 3’) and ctnnb2_ex16_whole R (5’ – GAGTTCTGTCTGATCGGGCC – 3’). PCR reaction was performed with Primestar HS DNA polymerase (Takara, cat # R10A) with the following settings: one cycle of initial denaturation at 98°C for 1 minutes, 30 cycles of denaturation at 98°C for 10 seconds, and extension at 68°C for 3 mins.

### Synthesis and cloning of 3’UTRs

For RT-PCR, total RNA was isolated from 1-8 cell wild-type and *ich* zebrafish embryos, using Zymo TRI Reagent (ZYMO Research, cat # R2050-1-50) as briefly follows: 20 embryos were collected in a 1.5 mL tube, and excess medium was removed. The samples were homogenized in 0.5 mL TRI Reagent, incubated for 5 minutes, mixed with 100 μL chloroform, and centrifuged at 10,000 × g for 15 minutes at 4°C. The aqueous phase was transferred to a new tube, mixed with 250 μL isopropanol, and incubated for 10 minutes. After centrifugation, the RNA pellet was washed with 70% ethanol, air-dried for 5–10 minutes, and dissolved in nuclease-free water.

Isolated total RNA was applied as template for cDNA synthesis, using the SuperScript IV UniPrime One-Step RT-PCR System (Invitrogen, cat # 12597025) according to manufacturer’s protocol. Wild-type RNA template was used for *ccna1, ctnnb2* and *tuba8l, ich* RNA for *ich-ctnnb2* with primers in Table 1.

Wild-type 3’UTRs were then cloned into pCS2+MT-AFP expression vector, where a S65A/Y145F mutated, humanized form of GFP, with improved fluorescent intensity is expressed from an SP6 promoter (Ogawa & Umesono, 2010). In the case of *ccna1* and *tuba8l* 3’UTRs, the DNA fragments were sticky-end ligated into *Xho*I and *Xba*I sites with T4 DNA Ligase (Thermo Scientific, cat # EL0014). Wild-type *ctnnb2* 3’UTR was blunt-end ligated into *Stu*I site in the same manner. All three ligations were then transformed into *E. coli* DH5a competent cells and selected on ampicillin containing LB plates. After inoculation and growth, miniprep was performed with GeneJET Plasmid Miniprep Kit (Thermo Scientific, cat # K0502). All plasmids were sent for sequencing for the verification of proper cloning. Probably due to the low quality of total RNA template, we were not able to amplify the whole *ctnnb2*^*ich*^ 3’UTR in one piece and hence we did the amplification in two steps as follows:

First, using ctnnb2_ex16-XbaI-R1 and ich-ctnnb2-F1 (Table 2) primers, we amplified a 1410 bp long fragment and blunt-end cloned into a *Eco*RV-linearized pBluescript KS(+) plasmid. ich-ctnnb2-F1 was designed to be specific to a region on the *ich* locus where it can have an ATC nucleotide triplet on its 5’ end. In this way, it is able to complete the residue of *Eco*RV recognition site on the pBluescript KS(+) plasmid, allowing us to perform another round of *Eco*RV digestion and blunt-end cloning of the second half of *ich-ctnnb2* 3’UTR (a 2114 bp long fragment amplified with ctnnb2_ex15-XhoI-F1 and ich-ctnnb2-R2 primers). After the completion of cloning, *ctnnb2*^*ich*^ 3’UTR containing pBluescript KS(+) was double-digested with *Xho*I and *Xba*I, which was followed by the gel isolation of *ctnnb2*^*ich*^ 3’UTR and sticky-end ligation into pCS2+ MT-AFP expression vector (as previously described).

**Table 2.**
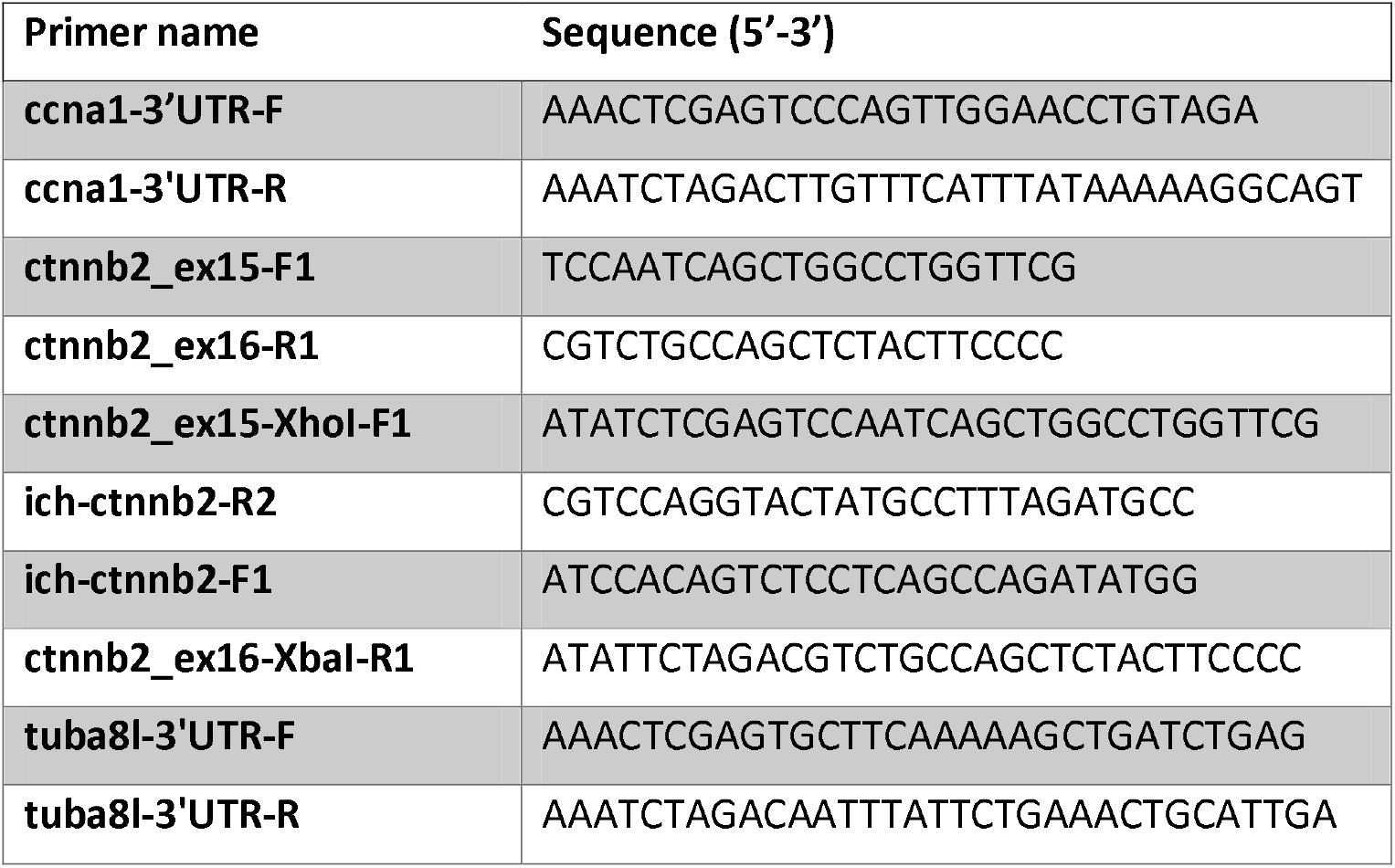
Primers used for 3’UTR amplifications. For *ccna1* and *tuba8l* 3’UTR, the primers were implemented from (Giraldez *et al*, 2006).

### mRNA synthesis

3’UTR containing pCS2+MT-AFP and unmodified pCS2+mCherry expression vectors were linearized with *Not*I and transcribed with mMESSAGE mMACHINE™ SP6 Transcription Kit (Invitrogen, cat # AM1340) following the manufacturer’s protocol. mRNA samples were purified with phenol:chloroform extraction and isopropanol precipitation method (according to the manufacturer’s protocol).

### Embryo injections and quantification of fluorescent signal

Wild type (*ekwill*) embryos were injected with 1 nL of the mixture of *AFP-3’UTR* and *mCherry* mRNAs at one cell stage. The concentration of *AFP-3’UTR* and *mCherry* mRNAs were 140 ng/μL and 100 ng/μL, respectively. At ∼24 hpf we documented AFP- and mCherry-expressing fish using a Zeiss Lumar.v12 UV stereomicroscope. For the evaluation of relative fluorescent protein brightness, we utilized Fiji (Schindelin *et al*, 2012).

### Screening of multiple *D. rerio* genomes for TE insertion

Transposable element sequences were retrieved from FishTEDB (Shao *et al*, 2024).

Multiple *D. rerio* genomes (Table 3) were screened for the presence of EnSpm-N49_DR, EnSpm-N49B_DR, and the hybrid of the two, EnSpm-N49/N49B_DR. The screening was performed using the BLAST search algorithm (Altschul *et al*, 1990) with default settings.

**Table 3.**
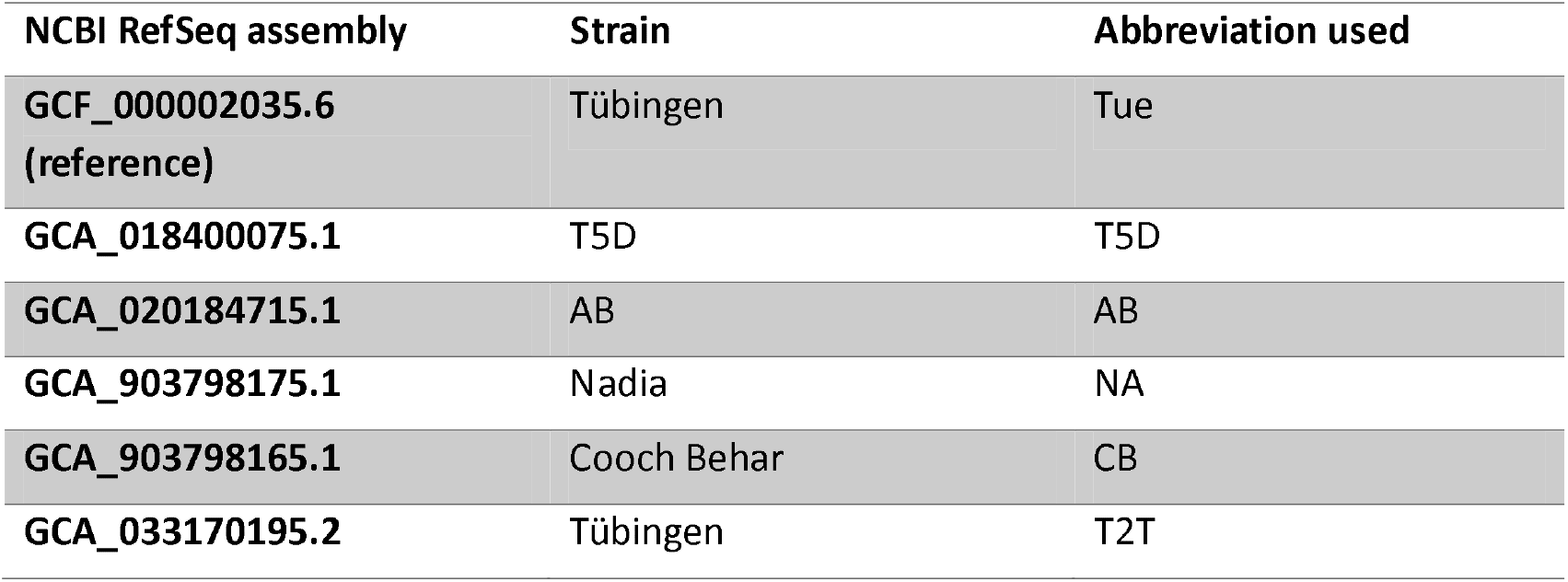
Genome assemblies screened for the presence of transposable elements.

| NCBI RefSeq assembly | Strain | Abbreviation used |
| --- | --- | --- |
| GCF_000002035.6<br>(reference) | Tübingen | Tue |
| GCA_018400075.1 | T5D | T5D |
| GCA_020184715.1 | AB | AB |
| GCA_903798175.1 | Nadia | NA |
| GCA_903798165.1 | Cooch Behar | CB |
| GCA_033170195.2 | Tübingen | T2T |

Screening results were further processed using custom scripts. TE hits were filtered downstream of the analysis for e-value of 1e^-10^. To alleviate spurious hits, a further filtering step was introduced whereby only hits where 80% of the transposable element was mappable were retained. All downstream analyses were performed on the hits retained after the filtering steps outlined above. Genome-wide insertion events were analyzed using *Granges* (Lawrence *et al*, 2013), and visualized using *circlize* (Gu *et al*, 2014). Annotation of the insertion sites was performed using *ChIPSeekR* (Yu *et al*, 2015; Wang *et al*, 2022).

### Statistics and visualization

Statistical analysis and visualization were performed in R (Team, 2018) using the *ggplot2* package (Wickham, 2016). All figures have been assembled in Affinity Designer (Serif Europe).

## Data availability

ONT sequencing data was uploaded in the Sequence Read Archive as BioProject PRJNA1207539. Telomere-to-telomere zebrafish genomic assembly can be accessed at BioProject PRJNA1029986. Code for the analysis of transposon positions is available on Github (https://github.com/danio-elte/zfish_transposon_screen). A frozen version of the repository with the raw numerical data retrieved from the analysis of fluorescence intensity and an annotated GenBank file of the *ctnnb2*^*ich*^ genomic sequence was uploaded to Zenodo as well (Varga *et al*, 2025).

## Author contribution

Conceptualization: SM, ESW, MV.

Data curation: MV, JO, SMB, FK.

Funding acquisition: MV, SMB.

Investigation: ZV, FK, SM, MV, ÁN.

Methodology: MV, JO, FK, ÁN.

Project administration and supervision: MV.

Writing – original and revised text: ZV, FK, ÁN, SMB, ESW, MV.

## Acknowledgements

We would like to thank Anita Rácz for fish care. This work was supported by the ELTE Eötvös Loránd University Institutional Excellence Program Grant 1783-3/2018/FEKUTSRAT. MV is a János Bolyai fellow of the Hungarian Academy of Sciences (BO/00555/22/8). This research was supported in part by the Intramural Research Program of the National Human Genome Research Institute (ZIAHG000183-24). The authors also acknowledge the support of the Freiburg Galaxy Team: Björn Grüning, Bioinformatics, University of Freiburg (Germany) funded by the German Federal Ministry of Education and Research BMBF grant 031 A538A de.NBI-RBC and the Ministry of Science, Research and the Arts Baden-Württemberg (MWK) within the framework of LIBIS/de.NBI Freiburg.

