## Supplemental Information for "Transposon insertion causes *ctnnb2* transcript instability that results in the maternal effect zebrafish *ichabod* (*ich*) mutation"

### Supplementary Information

**Supplementary Figure 1: Genotyping of the *ctnnb2* alleles.** (A) Schematic position of primers used for genotyping indicating the expected size of different amplicons. (B) Agarose gel showing the results of a genotyping assay in *ich* homozygous females. (C) Agarose gel showing the results of a genotyping assay in the embryos of an *ich^+/-^* incross.

**Supplementary Figure 2: Comparison of *ctnnb1* and *ctnnb2* sequence and expression.** (A) Schematic representation of Ctnnb1 (Uniprot ID: A0A5H1ZRJ2) and Ctnnb2 (Uniprot ID: Q8JID2) created with the *drawProtein* package (Brennan, 2018). (B) Overlay of Ctnnb1 and Ctnnb2 AlphaFold predicted structures using the pairwaise alignment tool of the RCSB Protein Data Bank (Jumper *et al*, 2021; Bittrich *et al*, 2024). (C) Dotplot of aligned *ctnnb1* and *ctnnb2* cDNA sequences created with *D-GENIES* (Cabanettes & Klopp, 2018). Results show extended similarities in the 5’UTR and CDS regions and little to no similarity in the middle of the 3’UTR. (D) Expression of *ctnnb1* and *ctnnb2* during oogenesis, based on single-cell analysis of zebrafish ovaries shows that both genes are expressed in post-meiotic cells and deposited to oocytes (perc_cell – percentage of cells showing expression; expr_lvl – expression level) (Liu *et al*, 2022).

**Supplementary Figure 3: Potential degradation of *ctnnb2^ich^* transcripts by the piRNA pathway.** (A) The structure of wild-type *ctnnb2* 3’UTRs as predicted by the ViennaRNA package (Lorenz *et al*, 2011) and SSRTool (Yang *et al*, 2022). (B) The stability of the 3’UTRs for the *ctnnb1, ctnnb2^wt^* and *ctnnb2^ich^* 3’UTRs as predicted by a random forest model (Vejnar *et al*, 2019), using 100 bp bins. Results suggest high stability for the *ctnnb1* 3’UTR and lower, stability for *ctnnb2* 3’UTR. Interestingly, this approach suggests that the sequence of the transposon itself has also high stability, so just stability considerations cannot account for the differences observed between *ctnnb2^wt^* and *ctnnb2^ich^*. (C) Developmental expression of EnSpm-N49 and EnSpm-N49B transposons based on Chang *et al*, 2022. (Note: the authors did not look for EnSpm-N49/N49B hybrid sequences specifically, but due to high sequence homology, a large fraction of EnSpm-N49B sequences are likely to be of hybrid origin in fact. Mean expression levels were calculated in the original paper.)

**Supplementary Figure 4: The characterization of EnSpm transposons observed in this study.** (A) Number of BLAST hits for EnSpm-N49 (orange), EnSpm-N49B (pink) and EnSpm-N49/N49B (green) transposon sequences, at different similarity values. The 80% threshold used in our study is shown by a dotted line. (B-D) Genomic distribution and number of the three different transposons. (E-H) Genomic positions of EnSpm-N49 (orange), EnSpm-N49B (pink) and EnSpm-N49/N49B (green) transposons in nonoverlapping 2-Mbp windows across the nuclear chromosomes of the indicated wild-type zebrafish strains.
